# Male competition and the evolution of mating and life history traits in experimental populations of *Aedes aegypti*

**DOI:** 10.1101/619072

**Authors:** Alima Qureshi, Andrew Aldersley, Brian Hollis, Alongkot Ponlawat, Lauren J. Cator

**Affiliations:** Department of Life Sciences, Imperial College London, Silwood Park, Ascot, UK SL5 7PY; School of Life Sciences, École Polytechnique Fédérale de Lausanne, Lausanne, Switzerland; Department of Entomology, Armed Forces Research Institute of Medical Sciences, Bangkok, Thailand, 10400

**Keywords:** (3-6)-*Aedes aegypti*, mating success, experimental evolution, male mosquito mating behaviour, reproductive control, sexual selection

## Abstract

*Aedes aegypti* is an important disease vector and a major target of reproductive control efforts. We manipulated the opportunity for sexual selection in populations of *Ae. aegypti* by controlling the number of males competing for a single female. Populations exposed to higher levels of male competition rapidly evolved higher male competitive mating success relative to populations evolved in the absence of competition, with an evolutionary response visible after only five generations. We also detected correlated evolution in other important mating and life history traits, such as acoustic signalling, fecundity and body size. Our results indicate that there is ample segregating variation for determinants of male mating competitiveness in wild populations and that increased male mating success trades-off with other important life history traits. The mating conditions imposed on laboratory-reared mosquitoes are likely a significant determinant of male mating success in populations destined for release.

## Introduction

The Yellow Fever mosquito, *Aedes aegypti*, is both an important vector of viruses and a main target of current reproductive control efforts. Mosquito reproductive control strategies [1] are in various stages of implementation [2–7], with facilities required to produce male mosquitoes at an industrial scale (millions/week) in order to sustain mass releases into the wild [8]. At this scale, even small deficits in the competitive mating success of released males against wild males potentially translate to large production and economic costs [9,10]. Release males will likely undergo many generations in the laboratory with potential for the mating competitiveness of these strains to be reduced by both a loss of heterozygosity [11] and laboratory selection [11,12]. A clearer understanding of the determinants of male mating success, and how these are affected by the rearing environment, is critical for optimizing mass-rearing and release strategies.

Mating in *Ae. aegypti* occurs in aerial swarms which are primarily composed of males with single females entering and being intercepted in much smaller numbers [13,14]. Males use the flight tones produced by females to detect potential mates in the swarm, responding to tones between 200-800 Hz with rapid phonotaxis [15,16]. If a male reaches a female, he must complete a series of mid-air manoeuvres to secure and perform insemination. The entire mating interaction lasts seconds and often the pair remains aloft for the duration [13,17]. During a mating attempt, females exhibit rejection behaviours in the form of kicks and leg thrusts that can effectively displace males through much of the mating interaction [15,17–20]. Recent work has suggested that acoustic interactions influence the outcome of mating attempts [19]. Opposite sex pairs of *Ae. aegypti* have been reported to actively adjust their flight tones to overlap at harmonic frequencies [21,22]. This phenomenon, termed harmonic convergence, has been found to be predictive of a successful mating attempt [19]. Converging males appear to offer no direct benefits to females or parental care to offspring but may offer indirect genetic benefits to females. The sons produced by converging pairs are more likely to exhibit both harmonic convergence and mating success [19]. While recent work has reported that males may also offer material benefits in the form of accessory gland proteins [23], it is not known whether females choose males based on variation in these benefits.

Therefore, the mating system of *Ae. aegypti* shares many characteristics with a lek [24], a group of displaying males which have aggregated for the sole purpose of mating [25]. As with other systems in which females choose mates based on genetic benefits, we would expect that over time female choice would erode genetic variation in male mating success. However, high variation in courtship signals in other animals has been reported to be maintained [26]. One possible solution to this “lek paradox” [27] is that additive genetic variation for male mating success is maintained because mating success is condition-dependent [28] and therefore correlated with other important life history traits which are under natural selection [29].

Experimental evolution is a powerful approach for investigating sexual selection [30]. Experimental evolution under different mating systems has been used in other insects to investigate the effect of sexual selection on components of nonsexual fitness [31,32], rates of adaptation [33,33–35], and the evolution of sexual traits including chemical [36] and acoustic [37] courtship signals. Here, we manipulated the opportunity for sexual selection in replicate populations of *Ae. aegypti* by controlling the number of males competing for a single female. We began our experiments with a large pool of eggs derived from a wild population. In one regime, mating occurred between isolated male-female pairs, eliminating male competition and female choice. In the other regime, five males competed to mate with a single female. We hypothesized that there was sufficient standing genetic variation in traits determining male mating success in a wild *Ae. aegypti* population to allow for an evolutionary response to the selection regimes within a limited number of generations.

In addition to harmonic convergence signalling and male mating competitiveness, several other traits such as body size [38] and sex ratio [39] have been reported to be heritable in *Ae. aegypti*. We also tested the hypothesis that the evolution of traits associated with male mating success would have correlated effects on other life history traits. We found that populations exposed to higher levels of male competition evolved higher competitive mating success relative to populations evolved in the absence of competition, with an evolutionary response visible after only five generations. There were also significant effects of the mating regime on the probability that a male is accepted by a female in isolated pair experiments, female body size and the number of eggs produced by females in the first clutch. This is the first time an experimental evolution approach has been applied to investigate sexual selection in mosquitoes. Our results indicate that there is ample segregating variation for male mating success in wild populations and that this variation trades-off with other important life history traits.

## Methods

### Maintenance of Evolved Populations

Experimental populations originated from collections of immature mosquitoes made from water storage containers (n=17) in two villages located in Muang District, Kamphaeng Phet Province (KPP), Thailand between February-April 2016. We collected 4500 eggs from these individuals to start the experiment.

We established six mosquito populations from these eggs, three that experienced high male competition every generation (HMC) and three that experienced no male competition (NMC). Each population consisted of one hundred breeding females, but the number of competing males varied between the two mating systems. For each HMC population, one hundred groups each with five males and one female were established. In the NMC populations, one hundred groups each with a single male and a single female were established (further details of mating procedures can be found in Supplemental Information). Logistical constraints limited us to manipulating the mating system of two experimental populations in a given day. We designated an HMC/NMC pair that was manipulated on the same day as a “replicate”. Thus, there were three replicate pairs of HMC/NMC populations (Fig. 1). After the 8 hr mating period, pooled females from each population were offered a bloodmeal. We controlled for the number of females contributing to subsequent generations by monitoring the insemination rate (Table S1) and fecundity (Table S2) in each population every generation. After five generations of experimental evolution, populations were reared under common garden conditions (Fig. 1) to control for parental effects [31,32,40]. For measurement of male mating competitiveness, mating and acoustic signalling in isolated pairs, and life history we used two experimental blocks. The blocks drew on the same set of eggs that were collected from the common garden rearing. All measures were taken over two blocks with the exception of female mating behaviour, which was measured in a single block.

**Figure 1.**
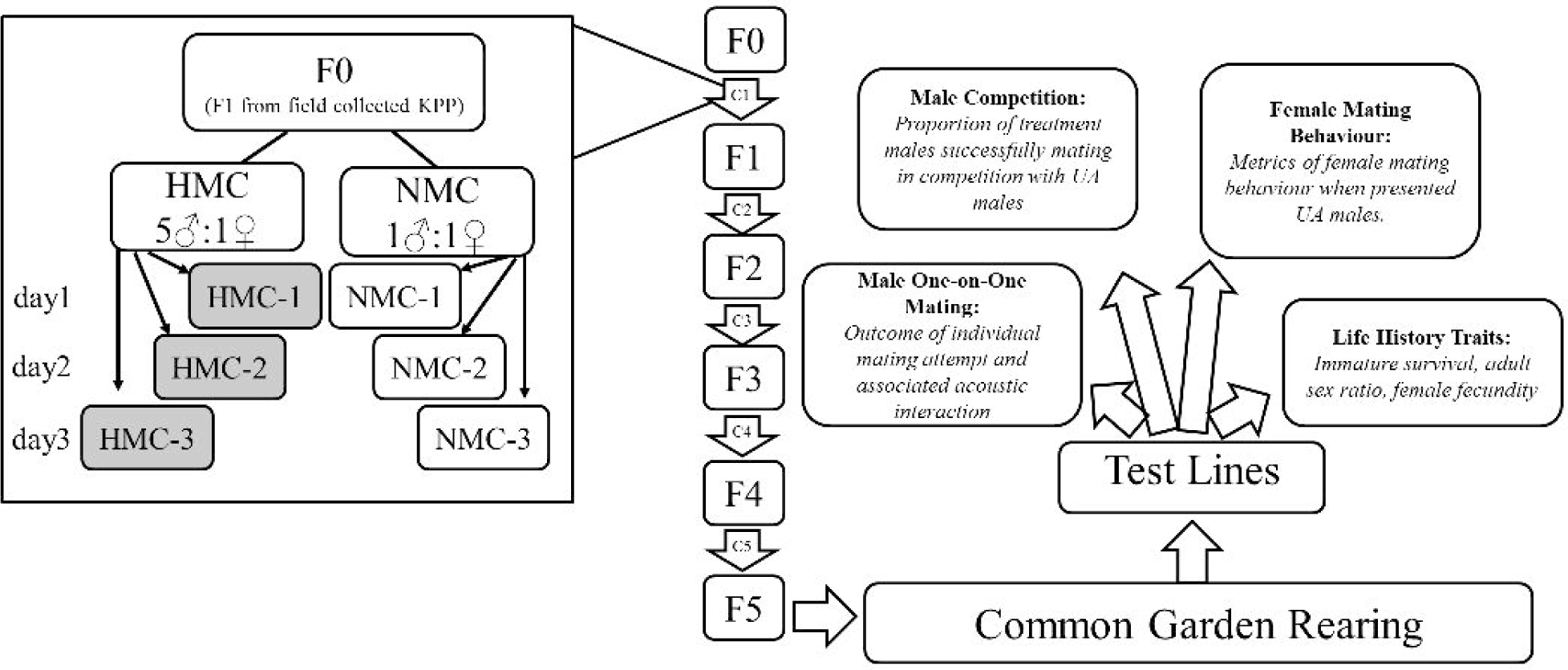
Overview of experiments conducted on selected populations. F0-F5 are the generations of the experimental evolution regime (these generated the HMC and NMC replicate populations), C1-C5 are the crosses that resulted in each generation. The inset shows an example shows how C1 resulted in

### Unselected Population

The unselected founding population (U) was created by rearing eggs taken from the original KPP pool under standard colony conditions (See Supplemental Information) for 2 generations to increase numbers. All U individuals used in phenotypic assays were F1-F3 from the field.

### Male Mating Competitiveness

Virgin males from each mating regime were lightly dusted [41] for identification. Two 2-5 day old virgin males from the U line and 2 males from either an HMC or NMC population were placed in a cage. An individual, 3-4 day old virgin U female was released into the cage and mating interactions were observed. Upon copula formation, we removed pairs and females were dissected to confirm insemination. We recorded dust colour and mating regime of males involved in each interaction. We ran 57-61 trials per experimental population over two blocks, discarding any trials in which no copula was formed or in which insemination status could not be determined.

### Mating and Acoustic Signalling in Isolated Pairs

The flight tones of paired mosquitoes were recorded as described in [17] (see Supplemental Information). For each pair, we analysed paired flight tone for the presence of harmonic convergence [21] and recorded whether the first attempted mating resulted in copula formation for a given pair. Harmonic convergence was defined as a matching of male and female harmonic frequencies during the mating attempt. Frequencies were considered to be matching if they were within less than 4.95 Hz and lasted a minimum of 1 s [19]. We ran 20-42 (HMC, n=100; NMC, n=124) of these trials per population over two blocks.

### Female Mating Behaviour

Virgin females collected from experimentally-evolved populations were released into mating cages containing four U males. For each trial, we recorded the total number of mating attempts, the time of each mating attempt, whether a copula was formed, and the start and stop time of the copula. We ran 30 of these mating trials per population, except HMC-2 which was not measured for this assay due to egg dessciation (HMC, n=60; NMC, n=90; U, n=90). Trials in which no attempts were made were discarded (15 out of 240 trials). We also measured the effect of mating regime on female fecundity. Females that were observed to form a copula were individually transferred to a modified 50 mL falcon tube. These females were then provided a bloodmeal and their first clutch of eggs was collected and counted. Females which did not mate, did not engorge when offered the bloodmeal, or died prior to laying their first clutch were excluded. The right wing of all females was removed and those that were not damaged were measured [42].

### Life History

Eggs from selected populations and the U line were hatched separately under a vacuum for 20 minutes and supplied with 0.1 mg of ground diet overnight. Larvae were separated into trays of 500 individuals in 2L of water and provided with 0.3 mg diet/larva/day. Each day we measured the number of living larvae in the trays and adjusted the amount of food provided. For each line, we recorded daily larval survival, daily pupation, daily adult emergence rates, and the sex ratios and body sizes of emerging adults. We recorded within-population individual fecundities for a subset of 10-30 females from each population after mating with males from the same population.

### Statistical Analyses

Unless otherwise stated all analyses were run in R version 3.1.1 [43] and the package “lme4”[44] was used to fit mixed models. We used the “afex” [45] package to run likelihood ratio tests and produce χ2 values and P-values for fixed effects. In all cases, replicate pair, replicate population, and (where appropriate) experimental block were incorporated as random effects. We describe the additional fixed effects for each model below.

We tested for an effect of mating regime (NMC/HMC) on the probability that a given male or female formed a copula using generalized linear mixed models (GLMM) with a binomial error distribution and logit link function. In the mating competition experiment we additionally tested the fixed effect of male dust colour (pink/yellow) and male wing length on the likelihood that a male successfully formed a copula in competition with U males. For isolated pair mating interactions, a GLMM fit with a binomial error distribution and logit link function was used to investigate the fixed effect of mating regime on whether harmonic convergence occurred during the interaction.

The effect of mating regime on female mating behaviours (attempt and copula latencies, total attempt durations, total attempt number, copula duration) was assessed using linear mixed models (LMM). We used a GLMM with a binomial response variable and logit link function to assess the effect of female mating regime on the probability of copula formation and sperm transfer to females.

A LMM was used to assess the fixed effects of female mating regime and wing length on fecundity. We made comparisons between mating regimes using a Sequential Bonferroni post-hoc test. We also determined the effect of mating regime on the proportion of first instar larvae that survived to emerge as adults using an LMM. The pattern of emergence over time was compared between mating regimes using a Mixed Effects Cox Regression to test for the effect using the ‘Survival’ package [46]. We used an LMM to test for the effect of mating regime, replicate, and block on the total proportion of emerging adults which were female. The winglengths of females and males from different mating regimes were compared using a LMM.

## Results

### Male Mating Competitiveness

We observed a total of 328 matings (HMC vs UA, n=147; NMC vs UA, n=181). HMC males were more likely to achieve a copula with a U female in competition with U males (56 ± 4.10%) than NMC males in competition with U males (40 ± 3.70%) (Fig. 2A, χ2=6.05, df_1_=1, P=0.01). The proportion of observed copulas in which males transferred sperm did not vary with male mating regime and insemination rates were generally high across replicates, populations, and mating regimes (78 ± 2.40%) (HMC♂, n=68, NMC♂, n=65, U♂, n=158; χ2=2.55. df_1_=2, P=0.28).

**Figure 2.**
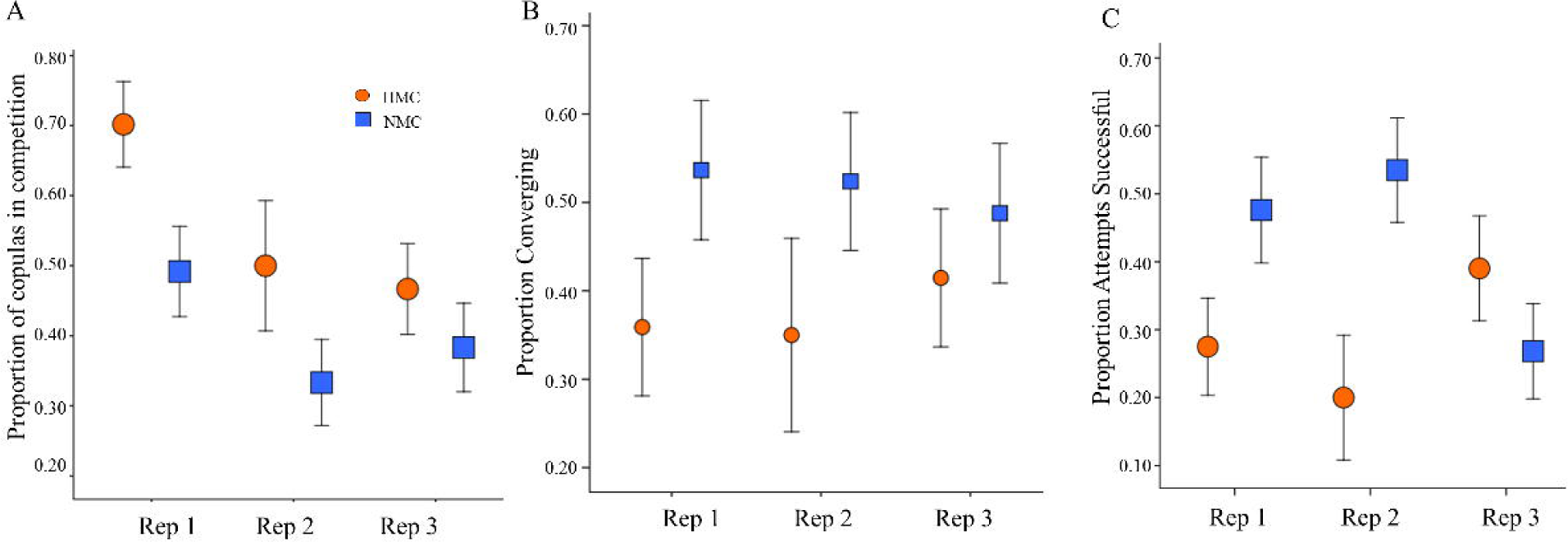
The effect of mating regime on male mating performance. A. Male mating success in competition with the unselected (U) line. Sample sizes, HMC-1=57, HMC-2=30, HMC-3=60, NMC-1=61, NMC-2=60, NMC-3=60 B. Proportion of mating attempts in isolated pairs containing a harmonic convergence event. C. Proportion of pairs in isolated pair mating experiments in which the first mating attempt was successful. Sample sizes for isolated pairs, HMC-1=39, HMC-2=20, HMC-3=41, NMC-1=41, NMC-2=42, NMC-3=41, error bars represent ± 1 SE.

### Mating and Acoustic Signalling in Isolated Pairs

We measured interactions between 224 isolated pairs (HMC vs UA, n=100; NMC vs U, n=124). Overall, 30.69 ± 4.61% of pairs formed a copula and 45.54 ± 3.33% converged at harmonics during the interaction. In these one-on-one mating attempts, NMC males were more likely to converge with a potential mate (Fig. 2B, χ2=4.16, df_1_=1, P=0.04). On average, NMC males were also more likely to successfully mate with the female in an attempt, but this effect of not significant when we control for population, replicate, and block effects (Fig. 2C, χ2=2.34,df_1_=1, P=0.13). As reported in previous work, the proportion of converging pairs was nominally higher amongst those that eventually formed a copula (41.18 ± 4.90% compared to 35.25 ± 4.34%). However, convergence did not significantly predict whether an attempt was successful (Figure S1, χ2=0.15, df_1_=1, P=0.70) and there was not a significant interaction between convergence and mating regime (Figure S1, χ2=2.07, df_1_=1, P=0.15).

### Female Mating Behaviour

We observed the responses of 225 females from evolved and unselected populations to unselected males (HMC, n=57; NMC, n=83; U, n=85). The total number of attempts (χ2=4.76, df_1_=2, P=0.09), latency to first attempt (χ2=0.07, df_1_=2, P=0.97), and attempt duration (χ2=0.41, df_1_=2, P=0.82) did not differ with female mating regime (Table S3). There was also no effect of female mating regime on the total rejections (χ2=2.12, df_1_=2, P=0.35), whether a copula was formed (χ2=0.99, df_1_=2, P=0.61), the latency between attempts starting and copula formation (χ2=0.02, df_1_=2, P=0.99), the duration of copula (χ2=0.60, df_1_=2, P=0.74) or whether a copula resulted in sperm transfer (χ2=0.21, df_1_=2, P=0.90).

Although there were no significant differences in female behaviour during the mating trials, we found differences between females from different mating regimes in reproductive output following the matings. Wing length did not significantly affect eggs laid in the first clutch (df=1, χ2=0.92, P=0.34). When we assessed the effect of mating regime removing the non-significant winglength covariate, there was a significant effect of female mating regime on the number of eggs laid in the first clutch (Figure 3A, S3, χ2=14.35, df_1_=2, P=0.0008). NMC females produced significantly more eggs (n=28; 57.57 ± 4.53 eggs/female) than U females (n=29; 38.41 ± 2.87 eggs/female) (Bonferoni Post-Hoc, P=0.04). While HMC females also produced more eggs (n=28; 52.21 ± 3.27 eggs/female) than U females, this difference was not significant (Bonferroni Post-Hoc, P=0.27).

**Figure 3.**
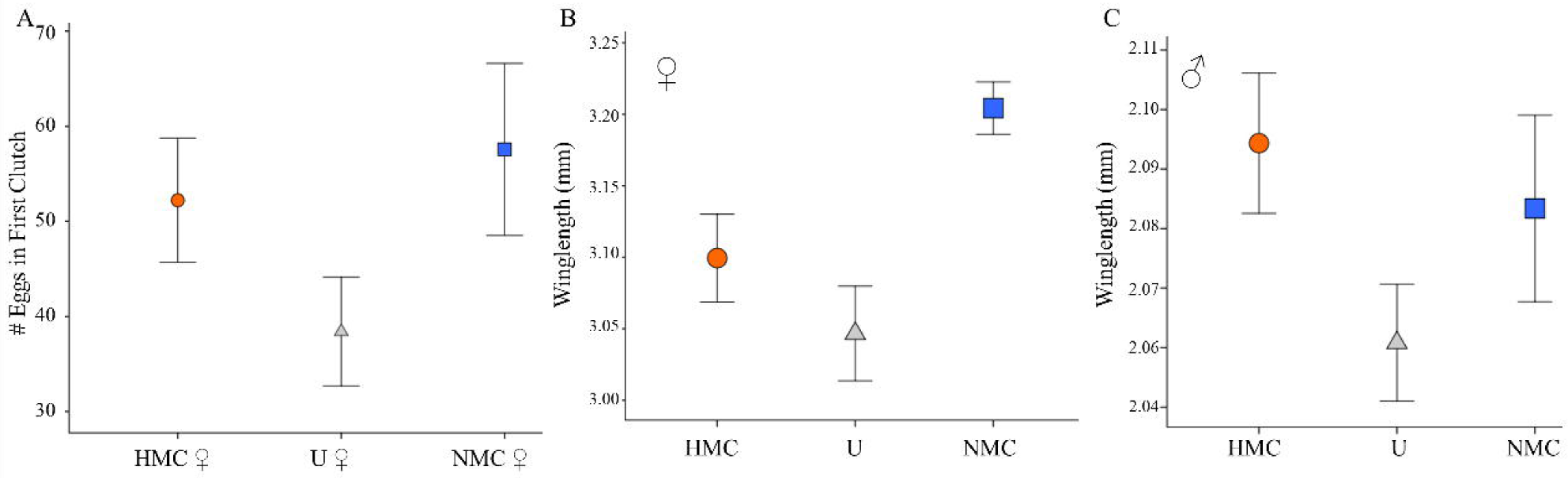
The effect of mating regime on female fecundity and male and female body size. A. The mean number of eggs produced by females mating with unselected males in female choice assays. Sample sizes, HMC=28 (HMC-1=16, HMC-3=12), NMC=28(NMC-1=12, NMC-2=4, NMC-3=12), U=29 (U1=11, U2=9, U3=9). B. The effect of mating regime on female body size. Samples sizes, HMC=35 (HMC-1=21, HMC-3=14), NMC=81 (NMC-1=24, NMC-2=31, NMC-3=26), U=39 (U1=11, U2=16, U3=12) C. The effect of mating regime on male body size. Sample sizes HMC=76 (HMC-1=36, HMC-2=14, HMC-3=26), NMC=68 (NMC-1=27, NMC-2=18, NMC-3=23), U=157 (U1=43, U2=49, U3=65). All error bars represent ± 1 SE.

### Effect of mating regime on life history traits

Mating regime did not significantly affect immature survival (Table 1, Fig. S2, χ2=1.67, df_1_=2, P=0.43) and there was no effect of mating regime on adult emergence time (Cox Regression χ2=1.25, P=0.21). The normal sex ratio in *Ae. aegypti* approximates 1:1, but may vary between populations [39,47,48]. Only the HMC populations produced significantly more males (∼60%, Table 1) than expected in a 1:1 ratio (χ ^2^ = 28.24, p < 0.0001). However, comparing between populations we found no significant differences in the proportion of males that emerged (χ2=5.61, df_1_=2, P=0.08). NMC females were significantly larger than those from the other two mating regimes (χ2=21.26, df_1_=2, P<0.001). There was no effect of mating regime on male body size (Fig. 3C, χ2=5.31, df_1_=2, P=0.07).

**Table 1.** Immature survival and sex ratios from mating regime and U populations. We report the mean ± 1 SE and * indicates a significant difference between the observed female:male ratio and that expected with a 1:1 female:male ratio.

| Mating regime | Replicate trays | Proportion of emerged adults that were female | Proportion of 1<sup>st</sup> instar larvae that became adults |
| --- | --- | --- | --- |
| U | 8 | 0.47 ± 0.01 | 0.82 ± 0.01 |
| HMC | 5 | 0.40 ± 0.02* | 0.71 ± 0.09 |
| NMC | 5 | 0.45 ± 0.02 | 0.65 ± 0.07 |

## Discussion

Most reproductive control programmes require the standardized rearing of mosquitoes at a large scale over many generations. Improved understanding of how evolution in the laboratory environment shapes mating traits could be used to mitigate negative effects of colonization and long-term rearing on release lines. In mosquitoes, long-term laboratory culture has been shown to lead to the evolution of increased testes size, earlier sexual maturation, and decreased sperm quality [11,12,49]. Here, for the first time, we directly assessed the effect of the selective environment on the evolution of male mating traits in *Ae. aegypti*. Our results provide strong evidence that in *Ae. aegypti*, changes in male mosquito mating phenotypes and other life history traits can evolve within only a few generations. This indicates that there is a relatively large amount of genetic variation for competitive mating success segregating in natural populations.

In particular, our results emphasize the importance of male competition for shaping mating success in competitive scenarios. Enforcing relatively high levels of male competition in mass-reared populations may be more conducive to maintaining traits contributing to male sexual success in the wild. While males from HMC populations won a greater proportion of matings in competition with U than males from NMC populations (Fig. 2A), they performed roughly equally to the U males overall, achieving only slightly more (56 ± 4.1%) of the total matings. Imposing high levels of sexual competition therefore maintained, rather than enhanced, performance in laboratory populations compared to the ancestral population. This suggests that mass-rearing operations would need to enforce very high levels of competition in order to maintain male competitiveness.

Production of other types of insects for mass releases have improved line quality by artificially selecting for competitive males [50–52]. In some of these instances, selection under competitive regimes has affected other aspects of life history such as fecundity [50,51], body size [31], and adult survival [31,50] over a similar number of generations to those in this experiment. A key advantage of experimental evolution is that it can provide information about genetic correlations and trade-offs between traits [30]. In our experiment, increased performance in competitive scenarios was correlated with evolved changes in other aspects of mating biology and life history.

First, we detected an apparent trade-off between competitive mating performance and harmonic convergence signalling. Manipulation of sexual selection in other insects has resulted in the evolution of signalling traits. For example, courtship song in *Drosophila pseudoobscura* was found to increase in intensity with increased sexual selection [53]. In *D. seratta*, cuticular hydrocarbon (CHC) profiles have been found to respond to sexual selection [54]. Here, we observed that NMC males were more likely to exhibit harmonic convergence during a mating attempt than HMC males (Figure 2B). Previous work has reported that the presence of harmonic convergence signals is correlated with female mating behaviours, suggesting that it may be used to inform female rejection responses [17,19]. In the absence of male competition, female rejection could have played a larger role in NMC male mating success compared to the HMC mating regime in which competition with other males for access may have been a more important factor. Thus, success in the NMC regime may have been more heavily dependent on male signalling and the associated female choice rather than male competition. Work in other species has provided examples of sexually selected traits that respond differently to male-male competition and female choice [55].

Alternatively, males from HMC populations may have evolved increased performance in other traits that enhanced success in competitive scenarios at the expense of traits underlying harmonic convergence ability. Two of the three HMC populations exhibited decreased performance in individual isolated mating attempts (Fig. 2C), further supporting the existence of trade-offs. Signalling dynamics may also differ in HMC versus NMC conditions such that acoustic dynamics vary with competitive scenario. We do not detect the established relationship between harmonic convergence and copula formation [17,18,21]. However, there is a trend in the data suggesting that the relationship between harmonic convergence and copula formation differs between the regimes (Fig. S1, Mating Regime x Convergence, χ2=2.04, df_1_=1, P=0.15). NMC populations nominally maintain the established positive correlation between the harmonic convergence and mating success, whereas HMC populations appear to lack this relationship. Future experiments could clarify this as well as identify which traits are most important for males achieving contact with the female in competition and which are most important for successfully mating with the female once access is gained.

The female mating behaviours we measured did not differ between evolved populations suggesting that the mating regimes did not select on female mating responses. Recent work in *Aedes* mosquitoes has demonstrated evolution of interspecific female choice over a similar number of generations [56,57]. Although our focus in this study was on pre-copulatory interactions, a number of post-copulatory changes in female mating behaviours [57–58] have been described in this species and these may have a larger role in sexual conflict. Future work could take advantage of an experimental evolution approach similar to our own to investigate the role of sexual selection in the evolution of female mosquito post-copulatory behaviour, as has been done in *D. melanogaster* studies focused on female sperm utilization refractory behaviours [58], reproductive output and timing [32,59,60], and resistance to male-induced harm caused by mating and seminal fluid proteins [59].

We observed evolved changes in other aspects of female life history. NMC females were larger than both U and HMC females (Figure 3B). Female body size in mosquitoes is a key determinant of bloodmeal size, which has important consequences for female reproduction and vectorial capacity [61–64]. There was no effect of regime on immature development so differences in adult female body size were not the result of differing developmental rates. We did not detect evolutionary change in male body size in our experiments, indicating that the increase in competitive pressure did not select for larger males.

Controlling for evolved differences between mating regimes in body size, NMC females laid more eggs in their first clutch than U females (Fig. 3A). In other dipterans, increased fecundity was reported to be related to serial selection [50] and populations from both mating regimes exhibited a nominal increase in fecundity relative to U females. However, only NMC populations exhibited a significant increase in fecundity compared to the U line (Fig. 3A). Further, this fecundity difference was only apparent when females were mated with U males (Fig. 3A). We did not detect any difference in fecundity when females mated with males from the same population (Table S2). This indicates that there may have been co-evolution between male and female traits determining female reproduction within the same line. Alternatively, because our experimental populations were also adapting to a novel laboratory environment, increased fecundity in these females could indicate that NMC females had adapted more quickly to reproduction in the lab environment which is known to select for early reproduction [52]. In other insects, there is conflicting evidence for the relationship between sexual selection and the nonsexual selection responsible for shaping adaptation to novel environments. In seed beetles, for example, sexual selection was found to accelerate adaptation to a novel environment (in this case a new seed host) [34]. However, other work in fruit flies suggested that populations evolving with stronger sexual selection do not adapt more quickly to thermal stress [33] or a novel diet [33,35]).

Differences between experimentally-evolving populations can sometimes arise due to differences in effective population size (N_e_). We mitigated any such effects by tracking female insemination rate (Table S1) and randomly selecting the same number eggs from every population each generation to ensure that a similar number of females contributed to the next generation. *Ae. aegypti* is thought to be monandrous and insemination rates were similar across regimes, suggesting that the number of males contributing to the next generation should have not differed systematically. Further, previous work has indicated that the effect of differences in effective population size in this kind of manipulation is minimal [24]. Just as relatively small population sizes were necessary in order to allow sufficient population-level replication in our experiment, we were similarly limited by practical constraints to handling our populations in replicate pairs instead of simultaneously. We incorporated this structure into our statistical modelling to control for differences between replicate pairs that arise from handling on one day versus another through the course of our assays. The mating regime differences we report here are therefore large enough to be detected despite any variation introduced by replicate pairs being manipulated on different days and replicate population. Our relatively simple selection regimes allowed us to manipulate sexual selection while equalizing other aspects of the mosquito life cycle, but these regimes do not reflect a natural situation and care should be taken when generalizing these results to selection in wild *Ae. aegypti.*

We chose to focus our assays on pre-copulatory mating behaviours and several key life history traits. Future work could expand on the traits measured to include post-copulatory competition and behaviour as well as additional life history parameters. A characterization of the traits involved in male mating success, and the extent to which manipulation of laboratory culture conditions affects these traits, would provide the basis for an evidence-based approach for how best to maintain these animals over successive generations in the lab in order to maximize production while maintaining male mating competitiveness. Future experiments could incorporate longer term selection and mesocosm work to increase our ability extrapolate the field. The experimental evolution framework demonstrated here offers a powerful new tool for further investigating sexual selection in mosquitoes.

## Ethics

Use of animal blood for maintenance of colonies was approved through Imperial College London’s Ethics Committee.

## Data

All data is available upon request

## Competing interests

We have no competing interests

## Authors contributions

LJC and BH conceived and designed experiments with input from AQ and AA. AP collected the wild population from KPP. All crosses were performed by AQ, LJC, and AA. All larval development and survival data was collected by AQ. All mating competition experiment were performed by LJC and AQ. All isolated pair experiments were conducted by AA. LJC analysed mating competition, isolated pair, and female behaviour data. AQ analysed all mosquito development and adult sex ratio data. LJC and BH lead the writing of the manuscript with inputs from, AA, AP and AQ. All authors gave final approval for publication.

### Acknowledgements

We would like to acknowledge Olivia Bates for providing pilot data on insemination rates used to choose the mating regime duration for crosses. We thank Laura Harrington and Garrett League for feedback on a late stage draft of the manuscript.

## Funding

This work was funded by the British Biotechnology Research Council Standard Award (BB/N003594/1) to LJC and AP and National Institutes of Health R21 Award (R21AI118593) to LJC.

