## Supplemental Materials for "Male competition and the evolution of mating and life history traits in experimental populations of *Aedes aegypti*"

**Additional Details of Methods**

**Maintenance of Evolved Populations.** Experimental populations originated from pupal and larval collections made from water storage containers (n=17) in two villages located in Muang District, Kamphaeng Phet Province (KPP), Thailand between February-April 2016. Emerging adults were identified to species and provided with a bloodmeal to produce F1 eggs. In order to induce hatching, a mix of 4500 F1 eggs was placed under a vacuum for 20 minutes. Newly emerged larvae were provided with 0.1 mg of ground fish food (Chiclid Gold [#04328], Hikari, Himeji City, Japan) and held in a 27 °C incubator overnight. First instar larvae were sorted into trays of 500 larvae/1L water and provided fish food pellets *ad libitum.* Individual pupae were placed into tubes plugged with cotton wool. Newly emerged adults were transferred using mouth aspirators to sex-specific cages, and provided a 10% sucrose solution for 3-7 days after emergence before being sorted into one of the two mating regimes.

For each cross, we set up individual matings so that each set of males and females experienced the regime in isolation. Mating arenas consisted of 22 x 8 x 16 cm plastic containers with an 18 x 6 cm mesh window in the lid. In preliminary experiments, mating readily occurred in these containers at both male-female ratios chosen. Males were observed to fly during the mating period in the stereotypical figure eight pattern associated with male swarming behaviour in the laboratory and field [1,2]. For crosses, virgin 3-7 day old males were selected at random and added to mating arenas. High male competition (HMC) populations were loaded with five males and No male competition (NMC) populations were loaded with a single male Males were left in arenas overnight to acclimatize. The mating arenas were supplied with 10% sugar solution. The following morning, between 0730-0900 we released individual females into mating arenas so that one female joined each arena. Mating regimes were imposed for a period of 8 hours. Females were removed with mouth aspirators from containers and transferred to population specific feeding cages. We set up 100 mating arenas per treatment and replicate population for a total of 600 crosses per generation. A subset of females were removed from each population to check insemination rates and body size.

To propagate the six populations, females were offered a bloodmeal of defibrinated horse blood (First Link Ltd. Wolverhampton, UK) using a membrane system (Hemotek Ltd., Blackburn, UK) (Approved by Imperial College London Health and Safety). Differences in fecundity between mating regimes may confound effects of selection and differential genetic drift caused by differing population sizes [3]. We controlled for the number of females contributing to subsequent generations by monitoring the insemination rate (Table S1) and fecundity (Table S2) in each population and generation. The number of females used in each generation was matched between all populations with a minimum of 80 females contributing.

Logistical constraints limited us to crossing two experimental populations in a day. We paired one NMC population with one HMC population and handled one pair per day (200 matings in a day) over three consecutive days for each cross (Fig. 1). The order of these pairs in the three day period was randomized across the generations. We designated a pair that experienced crosses on the same day as a “replicate”. Thus, there were three replicate pairs of HMC/NMC populations. Replicate effects were incorporated into statistical analyses to account for any effect that the day of crosses or subsequent bloodmeals may have had on populations (see Analyses section below).

**Unselected Population.** The unselected founding population (U) was created by rearing eggs taken from the original KPP pool under standard colony conditions. Larval rearing conditions were as described above and adults were held in mixed groups of up to 250 individuals with a 50:50 sex ratio (125♀:125♂) in 24 cm^3^ clear plastic cages. Rearing under these conditions took place over 2 generations to increase numbers. Thus, all U individuals used in phenotypic assays were F1-F3 from the field.

**Phenotypic measures.** Differences detected between traits in individuals produced by parents that experienced a particular sexually-selective regime may be due to evolutionary changes imposed by the regime itself or, alternatively, may be caused by the parental environment. For example, if females invest differentially in offspring under high versus no competition scenarios [4–6], we may observe variation in those offspring that do not reflect evolutionary change but rather a difference in parental investment (a non-genetic change). After five generations of selection, we subjected each population to one round of rearing under common garden conditions to eliminate potential differences driven by variation in parental environment (Fig 1). For common garden rearing, we used the standard larval conditions as described for the Life History assays described below. Adults were held in groups of 125 males and 125 females per line in 24 cm_­_^3^ clear plastic cages and provided with a 10% sucrose solution. After five days, mosquitoes were offered bloodmeals and the offspring collected from these individuals were used in phenotypic assays.

***Male Mating Competitiveness*.** Upon emergence males were transferred to population-specific cages and provided with a 10% sucrose solution for 2-5 days. Randomly selected males were cold anesthetized, placed onto petri dishes and lightly dusted [7] with pink or yellow dust (Swanda Inc. Stalybridge, UK). Colours were rotated between treatments for each day of experiments. For competition assays, two males from the U line and 2 males from either a HMC or NMC population were placed into 24 cm^3^ clear plastic cages. Males were provided with 10% sucrose solution allowed to recover and acclimatize overnight before proceeding. The following day, an individual 3-4 day old virgin U female was released into the cage and mating interactions were observed. Host odours were provided by the proximity of the experimenter. Each trial ran until the first copula was observed. Upon copula formation, pairs were removed, allowed to finish copulation and subsequently separated. Females were dissected to confirm insemination. We recorded dust colour and treatment of all males involved in each interaction. The right wing of both female and male were removed and measured as a proxy of body size [8].

***Mating and Acoustic Signalling in Isolated Pairs*.** The flight tones of paired mosquitoes were recorded as described in [9]. Briefly, individual 3-5 day old U virgin females were cold anaesthetized and tethered using glue (Nailene, Pacific World Inc., Aliso Viejo, CA, USA) to a short (~20mm) strand of human hair affixed to a metal pin. This was mounted on a stand in the centre of an 18cm^3^ white-mesh enclosure with an acrylic front face to allow for observation. The cage was set on top of a heating plate to provide temperature control throughout experiments, with host stimulation provided by experimenter proximity during interactions (<30cm) and by placing a worn sock between the recording chamber and the heating plate. Single 3-5 day old virgin males from one of the treatment populations were then released into the cage. For each pair, we analysed their acoustic signals for the presence of harmonic convergence [10] and recorded the outcome of the mating interaction. The possible outcomes were acceptance, in which a copula was successfully formed and males were able to clasp on to the female’s genitalia, and rejection, in which males were either kicked away or otherwise unable to form a copula (further description of harmonic convergence analyses and these behaviours can be found in [9]).

***Female Mating Behaviour*.** Upon emergence virgin females from each experimental population were transferred into population-specific cages and provided with a 10% sucrose solution for 2-4 days. Females were released into mating cages containing four 3-5 day old U males. Each trial ran for 4 minutes or until the first copula was observed. During this period we recorded the total number of mating attempts, the time of each mating attempt, whether a copula was formed, and the start and stop time of the copula.

We also measured the effect of mating regime on female fecundity. Females that were observed to form a copula with males from the U, HMC, or NMC line were individually transferred to a modified 50 mL falcon tube. These females were then provided a bloodmeal and their first clutch of eggs was collected. Any females which did not form a copula during the observations, did not engorge when offered the bloodmeal, or died prior to laying the first clutch were discarded from fecundity analyses. The right wing of all females was removed and undamaged wings were mounted onto a slide and measured as a proxy for body size.

***Life History*.** Eggs from selected populations and the U line were hatched separately under a vacuum for 20 minutes and supplied with 0.1 mg of ground diet overnight. Larvae were separated into trays of 500 individuals in 2L of water and provided with 0.3mg diet/larva/day. Each day we measured the number of living larvae in the trays and adjusted the amount of food provided. For each line, we recorded daily larval survival, daily pupation, daily adult emergence rates, and the sex ratios and body sizes of emerging adults. We recorded within-population individual fecundities for a subset of 10-30 females from each population after mating with males from the same group.

**Analyses**.

***Experimental Design.*** As described above, NMC and HMC populations were paired and designated as replicates, with one pair handled per day. There were 3 replicate NMC/HMC pairs. Male mating competitiveness, mating and acoustic signalling, and life history measurements were obtained in two blocks. A subset of eggs collected from the common garden rearing generation were hatched for each block so the individuals measured were all from the same generation. All replicates were run in each block and blocks were separated by 2 weeks. Again, replicate pairs were put through assays on successive days within the blocks. The HMC-2 population eggs desiccated between block 1 and block 2, so we were only able to obtain a single block of data for this population. Female mating behaviour measured in a single block and did not include the HMC-3 population. In the main text we report the sample size per treatment and replicate across the two blocks.

Unless otherwise stated all analyses were run in R version 3.1.1[11] and the package “lme4”[12] was used to run mixed models. We used the “afex”[13] package to run likelihood ratio tests and produce χ2 values and P-values for fixed effects.

***Male Mating Competitiveness*.** We used a generalized linear mixed effect model (GLMM) with a binomial distribution to test for the fixed effects of treatment (HMC/NMC), male dust colour (pink/yellow) and male wing length on the whether a population male formed a copula with the U female in competition with U males. We incorporated replicate pair, population, and block into our models as random effects. We used a similar approach to test for the effect of treatment on whether sperm was transferred. A GLMM with a binomial distribution was used to test for the fixed effect of treatment (HMC/NMC) on whether a mating resulted in successful sperm transfer. Replicate pair, population and block were again incorporated as random effects.

***Mating and Acoustic Signalling in Isolated Pairs*.** We used a GLMM with a binomial distribution to test for the fixed effect of treatment (HMC/NMC) on the whether harmonic convergence was detected during an isolated mating attempt (Yes/No) with a U female. In a separate model, we tested for the effect of convergence, treatment, and their interaction on whether a pair formed a copula (Yes/No). We incorporated replicate pair, population and block into our models as random effects.

***Female Mating Behaviour*.** The effect of mating regime and replicate on female mating behaviours (attempt and copula latencies, total attempt durations, total attempt number, copula duration) were assessed using linear mixed models (LMM). Female treatment was incorporated as a fixed effect and replicate as a random effect. We used a GLMM with a binomial response variable and logit link function to assess the effect of female mating regime as a fixed effect and replicate pair and population as random effects on the probability of copula formation and sperm transfer to females in these assay. A LMM was used to assess the effect of wing length and mating regime as fixed effect on the number of eggs laid.

***Life History*.** A GLMM was used to assess the effect of female treatment and wing length as fixed effects, and replicate pair and population as random effects, on fecundity. We made comparisons between mating regimes using a Sequential Bonferroni post-hoc test. We determined the effect of treatment on the proportion of first instar larvae that eventually emerged as adults using a GLMM with treatment as a fixed effect and replicate pair, population and experimental block as random effects. We used a GLMM to test for the effect of treatment on the total proportion of emerging adults which were female and incorporated replicate pair, population and block as random effects. The winglengths of females and males were compared using a LMM with treatment as a fixed effect and replicate pair, population and block as random effects. We made comparisons between mating regimes using a Sequential Bonferroni post-hoc test. The pattern of emergence over time was investigated using a Cox Regression in SPSS [14]. We tested for the effect of treatment on the probability of adult emergence controlling for replicate pair, population, and block.

**Supplemental Tables and Figures**

Table S1- Proportion of inseminated females from a subsample taken from each generation of crosses. Insemination was determined by examining the spermathecae for the presence of sperm. There were no significant differences in the proportion inseminated (Binary Logistic Regression, F1, P=0.99, F3, P=0.98, F5 P=0.85) in generations 1, 3 or 5 of the experiment.

| Generation | Treatment | N | Mean ± SE |
| --- | --- | --- | --- |
| F1 | H | 18 | 0.95 ± 0.01 |
|  | N | 28 | 0.79 ± 0.02 |
| F3 | H | 30 | 0.97 ± 0.01 |
|  | N | 18 | 0.73 ± 0.01 |
| F5 | H | 19 | 0.80 ± 0.02 |
|  | N | 29 | 0.67 ± 0.02 |

Table S2- Within line fecundity measured for first clutch. There were no significant differences between NMC and HMC lines (Bonferroni Post-Hoc, P<0.05).

| Treatment | Replicate | N | Number of eggs (Mean ± SE) |
| --- | --- | --- | --- |
| HMC | 1 | 4 | 27.50 ± 5.49 |
|  | 3 | 14 | 34.50 ± 4.02 |
|  | All | 19 | 39.84 ± 5.64 |
| NMC | 1 | 19 | 47.11 ± 3.40 |
|  | 2 | 5 | 45.40 ± 9.45 |
|  | 3 | 23 | 43.69 ± 3.32 |
|  | All | 38 | 38.61 ± 3.38 |

Table S3. Summary of mating interactions between unselected males and females from selected and unselected lines (mean ± SE). We also measured responses of unselected females to males as base line.

| Rep |  | Latency to First Attempt | Total Attempts | Total Attempt Period | Total Rejections | % Females forming Copula | Latency to Copula | Duration of Copula | %Sperm Transfer |
| --- | --- | --- | --- | --- | --- | --- | --- | --- | --- |
| 1 | NMC | 47.50 ± 10.72 | 1.37 ± 0.14 | 24.46 ± 9.45 | 0.67 ± 0.15 | 79 .00 ± 7.90 | 26.41 ± 8.90 | 12.55 ± 1.00 | 59.00 ± 10.70 |
|  | HMC | 34.51 ± 7.67 | 1.80 ± 0.28 | 40.17 ± 12.77 | 1.03 ± 0.29 | 82.76 ± 7.14 | 81.46 ± 13.86 | 14.33 ± 1.05 | 79.17 ± 8.45 |
|  | UA | 26.79 ± 4.91 | 1.67 ± 0.20 | 38.71 ± 11.09 | 0.97 ± 0.21 | 76.00 ± 8.10 | 33.64 ± 9.21 | 13.27 ± 0.94 | 59.00 ±10.70 |
| 2 | NMC | 43.35 ± 12.29 | 1.33 ± 0.18 | 36.23 ± 12.07 | 0.78 ± 0.16 | 70.00 ± 9.00 | 44.74 ± 13.21 | 15.79 ± 1.02 | 63 ± 11.40 |
|  | HMC | -- | -- | -- | -- | -- | -- | -- | -- |
|  | UA | 36.53 ± 7.06 | 1.73 ± 0.17 | 30.17 ± 8.14 | 0.99 ± 0.21 | 77.00 ± 7.90 | 30.57 ± 7.69 | 14.13 ± 0.82 | 78.00 ± 8.80 |
| 3 | NMC | 44.89 ± 8.95 | 1.40 ± 0.16 | 21.35 ± 7.76 | 0.93 ± 0.20 | 57.00 ± 9.50 | 25.59 ± 9.84 | 15.12 ± 1.50 | 88.00 ± 8.50 |
|  | HMC | 55.39 ± 10.64 | 1.63 ± 0.16 | 25.33 ± 10.00 | 1.07 ± 0.19 | 67.86 ± 8.99 | 61.45 ± 10.81 | 14.55 ± 0.97 | 63.16 ± 0.11 |
|  | UA | 74.73 ± 12.97 | 1.40 ± 0.17 | 22.28 ± 9.14 | 0.92 ± 0.16 | 69.00 ± 9.20 | 37.06 ± 13.39 | 14.28 ± 0.76 | 67.00 ± 11.40 |
| All | NMC | 45.30 ± 6.10 | 1.37 ± 0.10 | 27.28 ± 5.69 | 0.80 ± 0.10 | 69.00 ± 5.10 | 32.17 ± 6.23 | 14.36 ± 0.69 | 68.00 ± 6.20 |
|  | HMC | 44.77 ± 6.61 | 1.71 ± 0.16 | 33.02 ± 8.17 | 1.05 ± 0.17 | 75.44 ± 5.80 | 33.09 ± 8.36 | 14.43 ± 0.71 | 72.00 ± 6.90 |
|  | UA | 45.11 ± 5.43 | 1.60 ± 0.11 | 30.67 ± 5.48 | 0.95 ± 0.11 | 74.00 ± 4.80 | 33.49 ± 5.65 | 13.87 ± 0.49 | 68.00 ± 5.90 |

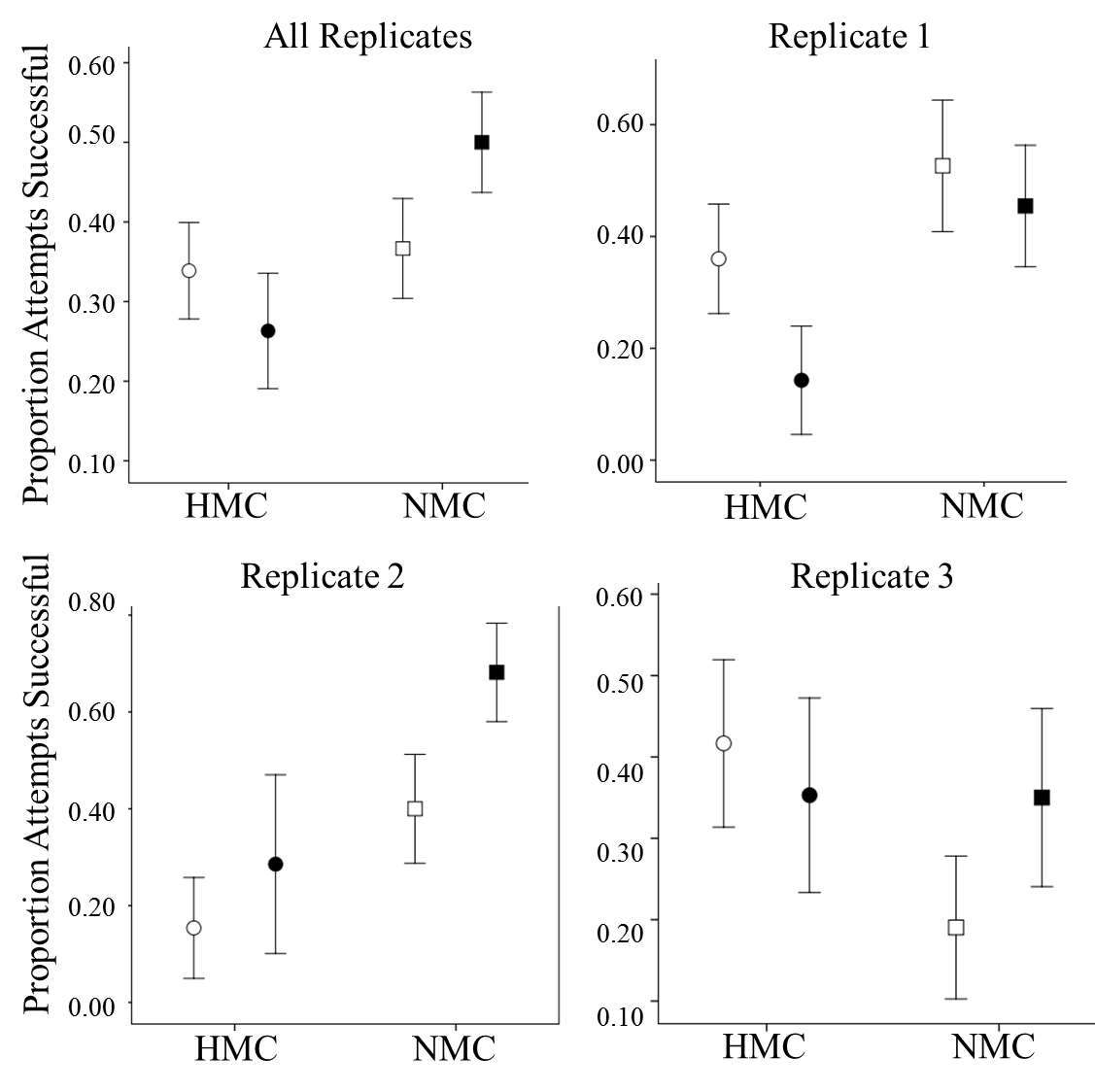

Figure S1. The effect of convergence on mating success. There is a trend suggesting that convergence only increases the mating success of NMC males (All Replicates). This trend is found in replicates 2 and 3, but not in replicate 1. None of the trends either at the replicate level or taking together all replicates are significant. Circles represent the HMC treatment, squares the NMC treatment. Darkened symbols indicate the presence of convergence in the mating attempt, unfilled indicate the absence of convergence during the attempt.

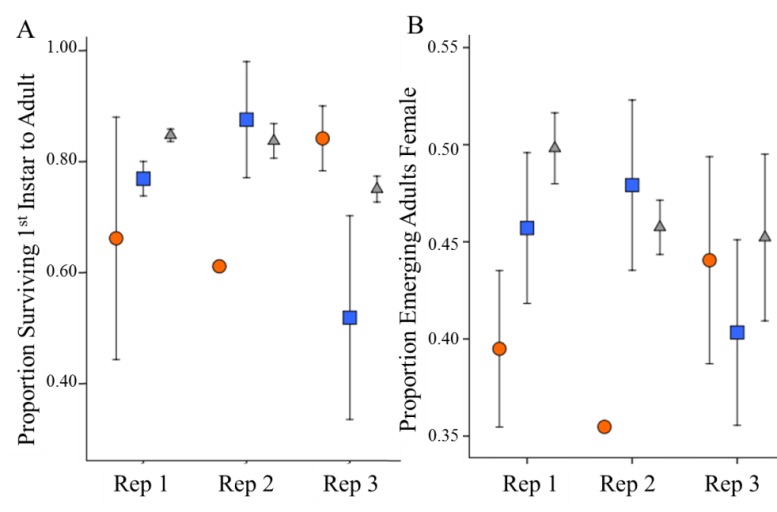

Figure S2. Life history data for each replicate. A. Proportion of 1^st^ instar larvae surviving to adulthood. B. Proportion of emerging adults which were female. HMC treatment=orange circles, NMC= blue squares, U=Unselected. Error bars represent 1 SE. Each mean is from two trays from the 2 experimental blocks. There was block for replicate 2 NMC. There were significant block, replicate, and treatment interactions.

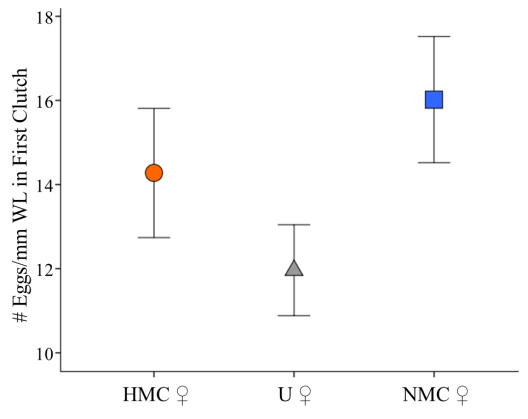

Figure S3. Eggs in first clutch per mm winglength for females from the three treatments. (n=55, HMC=15 (HMC-1=13, HMC-3=2), NMC=21(NMC-1=10, NMC-2=4, NMC-3=7), U=29 (U1=11, U2=7, U3=1)).

14. IBM Corp. Released 2016. IBM SPSS Statistics for Windows, Version 24.0. Armonk, NY: IBM Corp
